# A Novel Method for the Capture-based Purification of Whole Viral Native RNA Genomes

**DOI:** 10.1101/410282

**Authors:** Cedric Chih Shen Tan, Sebastian Maurer-Stroh, Yue Wan, October Michael Sessions, Paola Florez de Sessions

## Abstract

Current technologies for targeted characterization and manipulation of viral RNA primarily involve amplification or ultracentrifugation with isopycnic gradients of viral particles to decrease host RNA background. The former strategy is non-compatible for characterizing properties innate to RNA strands such as secondary structure, RNA-RNA interactions, and also for nanopore direct RNA sequencing involving the sequencing of native RNA strands. The latter strategy, ultracentrifugation, causes loss in genomic information due to its inability to retrieve unassembled viral RNA. To address this, we developed a novel application of current nucleic acid hybridization technologies for direct characterization of RNA. In particular, we modified a current enrichment protocol to capture whole viral native RNA genomes for downstream RNA assays to circumvent the abovementioned problems. This technique involves hybridization of biotinylated baits at 500 nucleotides (nt) intervals, stringent washes and release of free native RNA strands using DNase I treatment, with a turnaround time of about 6 h 15 min. RT-qPCR was used as the primary proof of concept that capture-based purification indeed removes host background. Subsequently, capture-based purification was applied to direct RNA sequencing as proof of concept that capture-based purification can be coupled with downstream RNA assays. We report that this protocol was able to successfully purify viral RNA by 561-791 fold. We also report that application of this protocol to direct RNA sequencing yielded a reduction in human host RNA background by 1580 fold, a 99.91% recovery of viral genome with at least 15x coverage, and a mean coverage across the genome of 120x. This report is, to the best of our knowledge, the first description of a capture-based purification method for assays that involve direct manipulation or characterisation of native RNA. This report also describes a successful application of capture-based purification as a direct RNA sequencing strategy that addresses certain limitations of current strategies in sequencing RNA viral genomes.

## INTRODUCTION

Viruses with RNA genomes are the cause of many infectious diseases with serious consequences to human health and mortality such as flaviviruses, HIV-1, SARS-coronavirus, HTLV-1 and influenza virus (Cantara et al. 2014). It has been shown that RNA-RNA interactions (Romero-López and Berzal-Herranz 2009) and intramolecular structure (Witteveldt et al. 2014) of viral RNA genomes play an important role in viral replication. However, extraction of viral RNA directly from culture often yields viral RNA with high host RNA background (Marston et al. 2013).

To deal with this issue, two main strategies are employed. Firstly, extracted viral RNA can be reverse transcribed to complementary DNA (cDNA) and amplified via polymerase chain reaction (PCR) to suit the sensitivities of the downstream assays (Ozsolak and Milos 2011). Following conversion to cDNA and amplification, *in vitro* transcription is then performed to retrieve genetic information in its RNA form, since both RNA-RNA interactions and RNA intramolecular structure can only be observed whilst in this form (Peattie 1979). Indeed, both Romero-López et al. (2018) and Witteveldt et al. (2013) amplified viral RNA obtained from cell culture via PCR amplification of viral cDNA clones followed by *in vitro* transcription. It has been shown, however, that such indirect methods for probing inherent properties of RNA increases experimental throughput at the expense of introducing PCR-induced artifacts and biases (Acinas et al. 2005), potentially clouding our understanding of these properties (Ozsolak et al. 2009). Secondly, isopycnic ultracentrifugation can be used to retrieve high purity samples of viral particles containing viral RNA, which can be subsequently extracted for downstream RNA assays. One example is the use of a sucrose density gradient to purify DENV2 RNA for RNA tertiary structure analysis (Dethoff et al. 2018). While the use of isopycnic centrifugation allows for a high purity of viral RNA, only RNA from encapsulated viruses are obtained (Liu et al. 2011), making this technique unfeasible for the study of unassembled viral genomes, which have also been shown to be crucial to our understanding of viral replication machinery (Manokaran et al. 2015; Moon et al. 2015).

With the advent of 3^rd^ generation nanopore sequencing, direct RNA sequencing of native RNA strands is heralded as a PCR and cDNA-bias free sequencing technology (Garalde et al. 2018). However, the direct RNA sequencing of RNA viral genomes is similarly affected by high amounts of host RNA background in viral cultures. The only method to date which addresses this issue is the use of custom sequencing adapters which has been demonstrated to yield high sequencing coverage for RNA viruses. Keller et al. (2018) utilized a custom sequencing adapter complimentary to the 3’ end of *influenza A* viral genome to selectively sequence only the viral RNA. However, this strategy is unviable for samples with diverse viral strains or genotypes where the 3’ sequence is unknown or non-conserved. An appropriate example is the wide diversity in *hepatitis C* virus genotypes, which has caused issues in PCR primer design for target enrichment strategies due to sequence ambiguity (Thomson et al. 2016). Similarly, Cowan et al. (2005) noted that PCR-based enrichment techniques where a priori knowledge of target sequences is required for PCR primer design render the enrichment strategies ineffective in the characterization of novel viruses. It follows that the same sequence ambiguity in viral genomes would pose a problem for using customized sequencing adapters during direct RNA sequencing.

From the abovementioned problems, we identified a need for an alternative technique that isolates viral RNA molecules from host RNA background without PCR amplification or cDNA synthesis. In the case of direct RNA sequencing, such a technique should address the limitations presented by current strategies. In this study, we describe a capture-based purification method that leverages on current nucleic acid hybridization and enrichment techniques. This novel method is able to isolate native viral RNA molecules from a high host RNA background without the use of PCR amplification or cDNA synthesis whilst retaining genetic information. Our capture-based purification method is distinguished from conventional hybridization-based or PCR-based enrichment techniques explored in the review paper by Mamanova et al. (2010) by virtue of not having any PCR amplification or cDNA conversion steps. Such a purification method, when merged with current technologies for probing and characterizing inherent RNA properties, could potentially catalyze the study of viral RNAs and elucidate previously inaccessible characteristics of viral RNA genomes.

Reverse-transcription quantitative polymerase chain reaction (RT-qPCR) was used as the primary proof of concept that capture-based purification indeed removes host background. Subsequently, capture-based purification was applied to direct RNA sequencing as proof of concept that capture-based purification can be coupled with downstream RNA assays. The application of capture-based purification to direct RNA sequencing as an alternative to current strategies will also be discussed.

## MATERIALS AND METHODS

### Viral culture

Viral inoculum and Huh-7 hepatocellular carcinoma cells (Japanese Tissue Repository) were cultured in DMEM media (10% Fetal Bovine Serum, 1% Penicillin Streptomycin) up to 70-90% confluency in Corning® 150 cm^2^ cell culture flasks (Sigma Aldrich). Prior to infection, spent media was discarded and Huh-7 cell monolayer was washed with 10 mL 1x PBS twice. Viral inoculum was provided by Low et al. (2006). A multiplicity of infection of 1 was used, 2 mL *dengue virus strain D1/SG/05K2402DK1/2005-1-I* (DENV1) was diluted with 2 mL pure DMEM media and mixed via pipetting up and down 5 times. Four milliliters of diluted virus was pipetted into each flask and incubated at 37°C for 1 h. Flasks were rocked every 15 min. Subsequently, inoculum was removed via pipette and 30 mL of pure DMEM media was pipetted to each flask was incubated at 37°C for 30 h.

### Trizol extraction of total RNA

Extractions were separately performed on viral supernatant and cell lysates. Media of infected flasks (viral supernatant) was transferred to a 50 mL Falcon tube. Subsequently the cell monolayer of each flask was treated with 8 mL of Trizol LS Reagent (Invitrogen) and 2.25 mL of 1x PBS (Thermo Fisher Scientific) followed by incubation at 37°C for 10 min. One milliliter aliquots of this cell lysate mixture were then pipetted to 1.5 mL Eppendorf tubes. 250 μL aliquots of viral supernatant were also pipetted into individual tubes and treated with 750 μL of Trizol LS Regeant. 200 μL of chloroform was then added to individual aliquots of viral supernatant and cell lysate before incubation at room temperature for 10 min. Tubes were shaken vigorously by hand for 30 s, and subsequently inverted by hand every 3 min. Each tube was centrifuged at 12 000 g, 4°C for 15 min. Upper aqueous phase in each tube was carefully removed and pipetted to a new 1.5 mL Eppendorf tube (approximately 500 μL). Two microliters of Glycoblue Coprecipitant (Invitrogen) and 500 μL of isopropanol (Sigma Aldrich) was added to each new tube and stored in −80°C overnight (or longer). Prior to the capture step, tubes were centrifuged at 12 000 g, 4°C for 15 min and supernatant was discarded. Pellet was re-suspended in 500 μL 75% ethanol diluted with nuclease-free water. Tubes were centrifuged again at 7 500 g, 4°C for 3 min and supernatant was discarded. Pellet was air dried. For the cell lysate RNA, total RNA concentration was determined via Nanodrop (Thermo Fisher Scientific) and re-precipitated in ethanol to obtain 30 μg total RNA pellets. Viral supernatant RNA either underwent capture or direct RNA sequencing library preparation without re-precipitation.

### Bait design

Twenty-one DENV1 baits were designed using BaitMaker (PLoS Neglected Tropical Diseases, Kamaraj et al. 2019, in press). The bait stock in our experiments was previously used for hybridization-based target enrichment in a published study by The Singapore Zika Study Group (Ho et al. 2017). Briefly, 120mer biotinylated DNA baits (Integrated DNA Technologies) specific to conserved regions across all serotypes of *dengue* virus were used. Baits were tiled along the viral genome with intervals of 500nt. A total of 6 pmol of baits were added to each capture.

### Viral RNA capture

This protocol was modified from that for the use of xGEN Lockdown baits and reagents (Integrated DNA Technologies), optimized for our purposes of RNA capture. SeqCap Hybridization and Wash Kit (Roche) was used in place of Integrated DNA Technologies reagents. Figure 1 demonstrates the workflow for the capture protocol.

**Figure 1:**
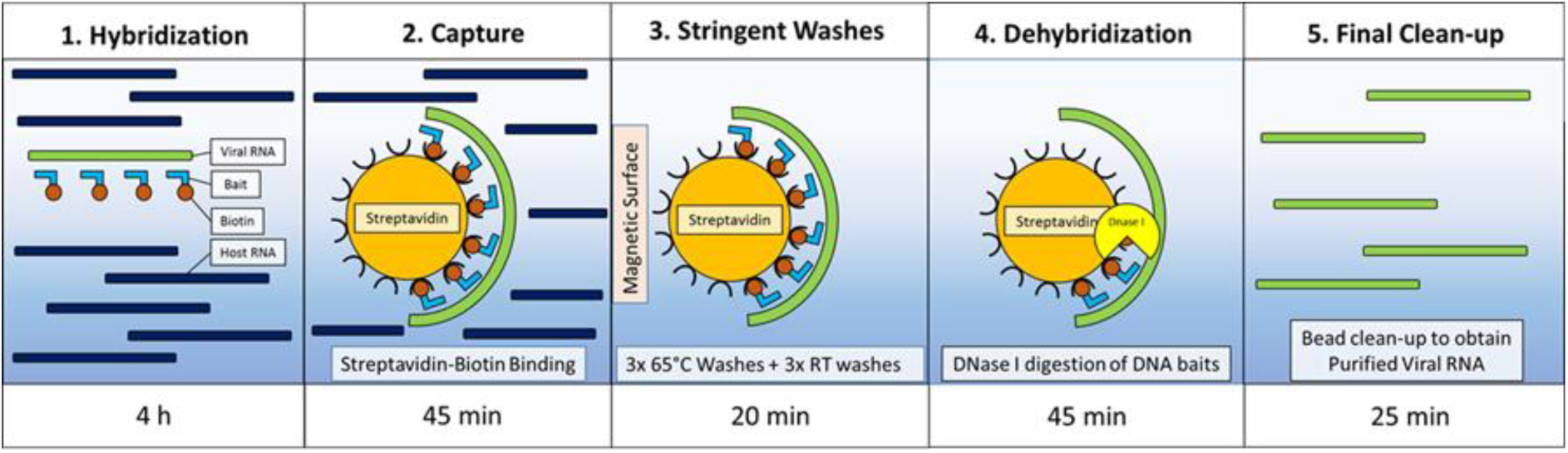
*Capture-based purification workflow*.

#### Hybridization

RNA Pellet from the extraction step was re-eluted with reagents shown in Table S1 and incubated at 65°C for 4 h in a thermocycler.

#### Streptavidin-Biotin Binding

100 μL of Dynabeads M-270 Streptavidin beads (Thermo Fisher Scientific) was pipetted to a 1.5 mL Eppendorf tube and equilibrated at room temperature for 30 min. The tube was put on a magnetic rack (Invitrogen) until eluate was clear and supernatant was discarded. 200 μL 1x Bead Wash Buffer was added and mixture was vortexed for 10 s before placing it on a magnetic rack and discarding the supernatant for a total of two washes. Hybridization mixture was immediately added to the beads and incubated at 65°C for 45 min.

#### Stringent Washes

All wash buffers were diluted to 1x prior to washing. Wash Buffer I and Stringent Wash Buffer was pre-heated for 1 hr to 65°C. 100 μL of Wash buffer I was added to the hybridization-bead mixture, agitated by hand and transferred to a new 1.5 mL Eppendorf tube. The new tube is placed on a magnetic rack. Once eluate is clear, supernatant was quickly discarded, taking caution not to remove any beads. 200 μL of Stringent Wash Buffer was added to the pellet and incubated at 65°C for 5 min, before discarding the supernatant as per the method above, for a total of two washes. Subsequent washes were performed as per the method above without the incubation step in the following sequence: Wash Buffer I, Wash Buffer II, and Wash Buffer III.

#### Dehybridization

Viral RNA was released from the beads via DNase I treatment. Beads were re-suspended in 20 μL of nuclease-free water, incubated at 95°C for 3 min then cooled to 37°C. Subsequently, 2.7 μL of 10x reaction buffer and 4 μL of TURBO DNase (Thermo Fisher Scientific) was added to the reaction and mixed by pipetting up and down 5 times, followed by further incubation at 37°C for 40 min. Subsequently, the tube was placed on a magnetic rack to allow the beads to aggregate. The supernatant was then transferred to a clean 1.5 mL Eppendorf tube for a 2.2x Agencourt RNAClean XP beads clean-up (Beckman Coulter). Briefly, 58.7 μL of beads were added to the supernatant and incubated for 8 min at room temperature, and then 2 min on a magnetic rack. The beads were washed twice with 75% Ethanol and residual ethanol was air-dried. The beads were then re-eluted in 30 μL of nuclease-free water.

### RT-qPCR

The proof of concept for our capture-based purification method was to demonstrate that host RNA background could be removed from target viral RNA. Absolute quantification of host and viral RNA was done using RT-qPCR. Two 1.5 ml Eppendorf tubes containing 30 μg of total RNA from cell lysate were prepared to represent pre and post-capture groups. 15 μL and 10 μL of nuclease-free water was added to the pre and post-capture group respectively. QuantiTect Probe RT-PCR Kit was used according to the manufacturer’s instructions. Primer targets were selected in the nonstructural 1 protein region of DENV1 and human GAPDH (glyceraldehyde 3-phosphate dehydrogenase) as a reference gene.

The volumes of reagents, shown in Table S2, were added to a 0.2 mL thin-walled PCR tube, and incubated in a ViiA 7 Real Time-PCR System (Applied Biosystems) with the program shown in Table S3 and analyzed using QuantStudio Real-Time PCR Software v1.1 (Applied Biosystems). Each plate included a non-template control for each pair of primers to check for PCR contamination. Samples were quantified in duplicate and non-template controls in triplicate.

A standard curve was constructed using a serial dilution of an oligo complementary to the primer target in accordance with MIQE Guidelines (Taylor et al. 2010). Custom oligoes of primer targets were ordered from Integrated DNA Technologies. Serial dilution was performed with nuclease-free water to obtain concentrations of 2.41×10^n^ copies/μL (where n=9,8,7,…,2). Each dilution was quantified in duplicate. The sequences of the oligoes, primers and probes used are shown in Table S4. Average C_q_ values for the serial dilutions were linearly regressed on log_2_(copies/μL) and Pearson’s product-moment correlation coefficient was calculated. Equation (1) was used to determine primer efficiency:

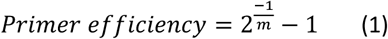

Where m is the gradient of the regression line of <C_q_> on log_2_(copies/μl)

Absolute quantity (copies/μL) was calculated using the respective equations of the regression lines.

The absolute quantity of DENV1 RNA, normalized to human GAPDH reference gene, was then compared between pre and post-capture groups. The ratio of the absolute quantity of DENV1 and human RNA pre and post-capture was compared, and purification factor calculated using Equation (2).

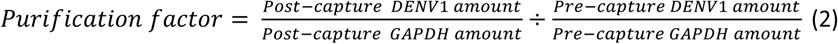

In this study we defined purification factor as a measure of how well host RNA background is removed from viral RNA.

### MinION Sequencing

Pre and post-capture RNA extracted from supernatant were sequenced on the MinION. Separately, we pooled 6 captures to generate the concentrated post-cpature group. Six captures each consisiting of 30uL of purified RNA was added to the same RNAeasy MiniKit (Qiagen) column and final elution volume was 30uL. Prior to library preparation, all RNA samples were polyadenylated using E. Coli Poly(A) Polymerase (New England Biolabs). The volumes of reagents shown in Table S5 were added to a thin-walled 0.5 mL PCR tube.

The reaction was then incubated in a thermocycler with the following program: 37°C for 30 min then cooled to 4°C before proceeding to a 2.2x Agencourt RNAClean XP cleanup (Beckman Coulter). The RNA was eluted in 11.5 μL nuclease-free water. Subsequently, library preparation and sequencing was performed according to the protocol for Direct RNA Sequencing Kit (SQK-RNA001) using 9.5 μL of RNA. All ligation reactions were extended to 20 min. The RNA Library was loaded on a R9.4 MinION flow cell and sequencing was ran for 48 h using MinKNOW v1.14.1 (GUI 2.1.14) interface. Live basecalling was turned off.

### Bioinformatic Analysis

Raw fast5 files were basecalled and end adapters were trimmed using Poreplex v0.2 (https://github.com/hyeshik/poreplex) which internally calls Albacore v2.3.1 (Oxford Nanopore Technologies). Basecalled read metrics and plots were then generated using NanoPlot v1.0.0 (https://github.com/wdecoster/NanoPlot). Subsequently, mapping and alignment was performed using Graphmap v0.5.2 (https://github.com/isovic/graphmap) with default parameters. Mean coverage and mapping quality was calculated using Qualimap v0.2.0 (García-Alcalde et al. 2012) respectively. Viral and human genome references used were EU081230.1 and GRCh38 respectively.

## RESULTS

### RT-qPCR

Based on the ratios of the absolute quantities of viral to host RNA shown in Table 1, we report a 561-791 purification factor, suggesting successful retrieval of viral RNA and substantial removal of host RNA background. Capture yield, which is defined as the percentage of viral RNA amount captured, was calculated to be 6.98%-40.4%.

Figure S1 shows the standard curves for DENV1 and GAPDH primers. The primer efficiency of DENV1 and GAPDH was 0.66 and 0.92 respectively. The DENV1 primer set used was a pre-optimized primer set that is able to detect all 4 *dengue* virus serotypes previously used and therefore contains ambiguous bases, explaining the lower primer efficiency. It was noted that relative quantification using the 2^-ΔΔCT^ method (Schmittgen and Livak 2008) was not appropriate as the primer efficiencies of DENV1 and GAPDH primer sets were not comparable. Absolute quantification and subsequent ratio comparison of viral and host RNA amounts is therefore more accurate in the determination of purification factor. All obtained values were within the dynamic linear range and limit of detection of the reaction conditions and primer sets used, suggesting reliability and accuracy of the reported RT-qPCR results.

### MinION sequencing

Based on the mapping rates to human and DENV1 reference genomes for pre and post-capture groups, shown in Table 2, purification factor was calculated to be 272 fold. We also report a 3.89 fold increase in percentage recovery of DENV1 genome over 15x coverage from a single pre-capture to a single post-capture RNA sample. On the other hand, comparing the concentrated post-capture group (containing a pool of 6 captures) to the pre-capture group yielded a purification factor of 1580 fold and a 14 fold increase in the percentage recovery of DENV1 genome above with at least 15x coverage from 7.05% to 99.91%. *Supplementary figures 2, 3, and 4* show representative coverage plots for the pre-capture, post-capture and concentrated post-capture groups respectively.

In Equation (3) and (4), a binomial distribution was used to determine the minimum read coverage, *n*, required to have a 0.95 confidence that *k* reads mapping to a particular base, *X*, are correctly called. We used the average error read rate of our runs as the probability, *p*, of correctly calling and mapping a single base.

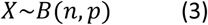

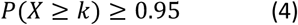

Since the common heuristic for variant calling with Illumina reads requires 10x coverage (Illumina, https://www.illumina.com/Documents/products/technotes/technote_snp_caller_sequencing.pdf), we assumed the criterion that at least 10 reads were correctly called and mapped to a single base (*k*=10).

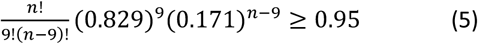

Based on our sequencing runs, the average error read rate of pre and post-capture sequencing runs was 17.1±0.8% (calculated from Table 2). With this error read rate, we substitute 1-*p*=0.171 into Equation (5) and determined that the minimum coverage, *n*, required for variant calling to be at least 15x.

## DISCUSSION

### Capture-based purification removes high host background

We demonstrated with RT-qPCR that our novel RNA capture technique can be used to retrieve and purify viral RNA in its native form for downstream assays. This was corroborated by our direct RNA sequencing results. Indeed, unpurified viral RNA represented by the pre-capture group did not produce sufficient throughput for any meaningful downstream analysis due to the indiscriminate nature of the sequencing of both host and viral RNA. However, after treatment with our capture method, the amount of host RNA background has decreased greatly as seen from the 272 fold increase in mapping rates to target viral RNA. In comparison, even with the incorporation of PCR amplification for shotgun sequencing of *enterovirus*, Jensen et al. (2015) reported only a 2-13 fold increase in mapping rates to *enterovirus*. Moreover, Wongsurawat et al. (2019) observed only a 36 fold increase in mapping rates to Zika virus (ZIKV). This suggests that our capture-based purification method is highly successful in the removal of host RNA background. The minimum coverage required for variant calling was, as described above, benchmarked to that of Illumina reads so that the effectiveness of our capture-based purification method could be more accurately evaluated based on the higher error read rates of direct RNA sequencing technology. Even with the higher adjusted heuristic for variant calling of 15x, the 3.89 fold increase in percentage of the genome above 15x coverage suggests that application of our capture-based purification method increases the amount of target genetic information retrieved.

### Capture-based purification is robust

Our method fundamentally allows for the capture of unknown viral strains and genotypes due to the flexibility of adding multiple baits complementary to different regions of the viral genome, or even multiple baits specific to different viruses in a single capture. Indeed, Deviatkin et al. (2017) has reported the successful use of genus-specific degenerate baits for hybridization-based enrichment, which can be seamlessly incorporated into our bait design. By extension, the baits used can be appropriately modified according to the intended research focus. The robustness of this capture-based purification method therefore not only broadens the applicability of direct RNA sequencing on studying unknown viruses but also the applicability of a diverse range of direct RNA probing technologies.

Another major advantage to our capture-based purification method is that large machinery like an ultracentrifuge is not required. To perform our capture method, only a thermocycler and a heat block is required. This is especially applicable to research environments where an ultracentrifuge is not readily available. More fundamentally, our capture-based purification method allows for the capture of both unassembled and assembled viruses, which any method leveraging on the differential physiochemical properties of encapsulated viral RNAs cannot. This would be important in studies of unassembled RNA segments, which have been demonstrated to be of great biological importance. Indeed, in the case of DENV1, subgenomic flaviviral particles have been shown to underlie mosquito immunity and viral transmission (Pompon et al. 2017). If so, a method which can remove host background and capture unassembled viruses would be useful for characterising inherent RNA properties of unassembled viruses.

### Limitations

One point of concern for our protocol is the low capture yield. However, it must be noted that the capture yield is still significantly higher than that of conventional viral purification methods. Hall et al. (2014), in a study evaluating conventional viral purification methods, reported a 100 fold decrease in viral copy number post-treatment, whereas our method results in, at most, a 14 fold decrease in viral copy number. In evaluating our capture-based purification method, a wide range in capture yield was observed, which was due to the high sensitivity of RNA to potential contamination during handling. This is a problem common to any viral RNA purification method.

Our capture-based purification method has a turnaround time of approximately 6 h 15 min (Figure 1), comparable to the time required for isopycnic gradient-based methods (Buclez et al. 2016). While there is a significant loss of viral RNA post capture, we also note that our capture method is scalable. Multiple captures can be performed and purified RNA pooled together before re-concentrating in the final bead clean-up step. The concentrated post-capture group was a demonstration of how our capture-based purification method was appropriately scaled to suit the sensitivity of direct RNA sequencing. Indeed, after comparison of the post-capture and concentrated post-capture sequencing runs (Table 2), the 2.5 fold increase in the percentage of reads mapping to DENV1 suggests that scaling our method greatly improved the signal-to-noise ratio of this particular downstream RNA assay. While it was opined that this improvement would similarly be observed in other downstream RNA assays, further experimental verification is required.

### Application of capture-based purification to direct RNA sequencing

The use of direct RNA sequencing to study viral genomes is limited by its sensitivity. Conventionally, the sequencing adapter used in direct RNA sequencing was a poly(T) adapter which would hybridize with poly(A) tails, facilitating adapter ligation (Garalde et al. 2018). This poly(T) adapter is also the sequencing adapter provided in the direct RNA sequencing library preparation kit (SQK-RNA001). Currently, four strategies may be employed for direct RNA sequencing of RNA viruses. Firstly, as described by Keller et al. (2018), a custom adapter can be used during library preparation. The second strategy is the use of the conventional poly(T) adapter supplied in the kit by polyadenylation of RNA prior to library preparation and sequencing. The third strategy is to couple polyadenylation with capture-based purification. The fourth is to couple polyadenylation with host rRNA depletion (Wongsurawat et al. 2019). A summary of the advantages and disadvantages for these strategies are presented in Table 3.

The custom adapter strategy employed by Keller et al. (2018) greatly improves sequencing specificity, and was demonstrated to yield 98.8% of reads mapping to *influenza A*. This strategy allows sequencing of RNA viruses with or without an inherent poly(A) tail since the conventional poly(T) adapter was substituted with a custom sequencing adapter specific to the target. However, one drawback of this strategy is that foreknowledge of the 3’ end of RNA targets is required. This entails that sequencing of unknown, novel, or mutant viruses would be difficult. Another limitation is that this strategy may be ineffective in cases where the RNA sample is fragmented since the 3’ end of each fragment of viral RNA is distinct.

Degenerate adapters may be used to circumvent the first limitation, as with the degenerate adapters at the +4 position used by Keller et al. (2018). However, using degenerate adapters would only be able to address the problem of mutant viruses but not that of novel or unknown viruses. Indeed, foreknowledge of target 3’ sequences is still required. On the other hand, attempts at using multiple adapters at different locations on the target genome cannot address the problem of RNA fragmentation. Custom adapters must be specific to the 3’ end of each fragment for ligation. In most laboratory samples, however, RNA is stochastically fragmented. As such, determination of the exact 3’ end of each fragment would be impossible and so such methods to circumvent the problem of RNA fragmentation are unviable.

In this study, we demonstrated the second and third strategies, which make use of the conventional poly(T) adapter. This was done using DENV1, a positive single stranded RNA virus lacking a poly(A) tail. The protocol used to generate the pre-capture group sequencing results employ the second strategy while that for the post-capture and concentrated post-capture results employ the third. The polyadenylation of RNA and subsequent use of the conventional poly(T) adapter has several advantages. First, the polyadenylation of RNA prior to library preparation and sequencing allows for sequencing of RNA viruses with or without an inherent poly(A) tail. Second, usage of the conventional poly(T) adapter removes the need for any foreknowledge of the target sequence. Third, all RNA fragments would be ligated with adapters since the adapters are not sequence specific, circumventing the problem of fragmented RNA. However, the main disadvantage strategies involving polyadenylation of RNA is that sequencing specificity is greatly reduced as host mRNA transcripts include a poly(A) tail and will also be sequenced. Wongsurawat et al. (2019) independently replicated our pre-capture group results but with ZIKV, which is also a positive single stranded RNA virus lacking a poly(A) tail in the family Flaviviridae. The authors observed 0.11% of reads mapping to ZIKV, which corroborates the low 0.63% of reads mapping to DENV1 in our pre-capture sequencing run. The authors also reported 3.89% of reads mapping to ZIKV whilst employing the fourth strategy. The low percentage of target reads underscores this main disadvantage of sequencing specificity.

The issue of low sequencing specificity is, however, less pronounced in the third strategy. Comparing the second strategy (pre-capture) and third strategy (post-capture), there is a 50 fold increase in percentage of reads mapping to DENV1. Furthermore, a comparison of pre-capture and concentrated post-capture results yields a 123 fold increase in mapping rates to DENV1. This suggests that while simply polyadenylating RNA samples leads to low sequencing specificity, applying our capture-based purification prior allows us to reap the advantages of using the conventional poly(T) adapter whilst having a significantly higher sequence specificity. Additionally, coupling capture-based purification with polyadenylation yields a higher target mapping rate of 31.31% with regard to the 3.98% observed when the fourth strategy was employed (Wongsurawat et al. 2019). This suggests that the third strategy is superior with respect to sequencing specificity when compared to the fourth strategy. It is noted, however, that the sequencing specificity of the third strategy was still lower than that of the first strategy. Moreover, applying capture-based purification in the third strategy significantly increases the turnaround time as compared to the first, second and fourth strategy. Therefore, it is suggested that the third strategy should only be employed where foreknowledge of the viral sequence at hand is limited, or where RNA integrity is low. It is also suggested that the second and fourth strategy be used in cases where detection rather than characterisation of RNA viruses is required due to significantly lower sequencing specificity.

In this study, we demonstrated a novel capture-based method for the purification of viral RNA from host RNA. In addition, we described a successful attempt of applying our capture method to the direct RNA sequencing of viral culture, using a DENV1 model, exemplifying how our capture method could significantly remove host background in downstream assays requiring viral RNA in its native form. Further improvements to the capture yield are underway and could potentially expand its application to clinical samples where starting RNA material is low. The fundamental advantages of using our capture-based purification method makes it a superior alternative to conventional viral purification methods and addresses limitations of current strategies for direct RNA sequencing, potentially paving a new path for the characterization and direct RNA sequencing of viral RNA.

## Supporting information

All supplementary tables and figures

All tables

## DECLARATIONS

### Authors’ contributions

CSC Tan was the primary contributor to this study with guidance from his mentor PF de Sessions and helpful advice from OM Sessions, S Maurer-Stroh and Y Wan.

## Acknowledgements

We would like to acknowledge Ng Hui Qi Amanda and Niranjan Nagarajan from Computational Systems Biology 5, GIS; Aw Poh Kim Pauline (DENV1 primer design), Kwan Kam Weng Philip, Balamurugan Periaswamy, Png Eileen from GERMS Platform for their technical advice throughout the course of this study; Vithiagaran Gunalan and Dimitar Kenanov from Maurer-Stroh’s group for helpful conversations. We would also like to acknowledge members of Wan Yue’s team, in particular, Lim Wei Sheng Shaun, Aw Jong Ghut Ashley. Lastly, we would like to thank Andreas Wilm and Shih Chih Chuan from the GIS Bioinformatics Core.

## Availability of data and materials

Data will be made available through publication and NCBI.

## Compliance with ethical standards

This article does not contain any studies with human participants or animals performed by the author.

## Consent for publication

All authors listed on this manuscript have read and agreed to the publication of this research.

## FIGURE LEGENDS

Table 1: Comparison of absolute quantities of DENV1 and GAPDH used for calculation of purification factor.

Table 2: Summary of bioinformatic analysis for pre and post-capture MinION sequencing runs.

Table 3: Summary of advantages and disadvantages of strategies for direct RNA sequencing of RNA viruses.

Table S1: Reagents and volumes for hybridization step.

Table S2: Reagents and volumes for RT-qPCR.

Table S3: RT-qPCR program set on Applied Biosystems ViiA 7 Real Time-PCR System.

Table S4: Primers, probes and standards used with the respective modifications for RT-qPCR.

Table S5: Reagents and volumes used for polyadenylation of RNA samples.

Figure S1: DENV1 and GAPDH standard curves used for calculation of primer efficiency.

Figure S2: Coverage plot against nucleotide position on DENV1 reference for pre-capture MinION sequencing run.

Figure S3: Coverage plot against nucleotide position on DENV1 reference for post-capture MinION sequencing run.

Figure S4: Coverage plot against nucleotide position on DENV1 reference for concentrated post-capture MinION sequencing run.

