## Supplementary material for "A Novel Method for the Capture-based Purification of Whole Viral Native RNA Genomes": All supplementary tables and figures

Figure 1: Capture-based purification workflow.

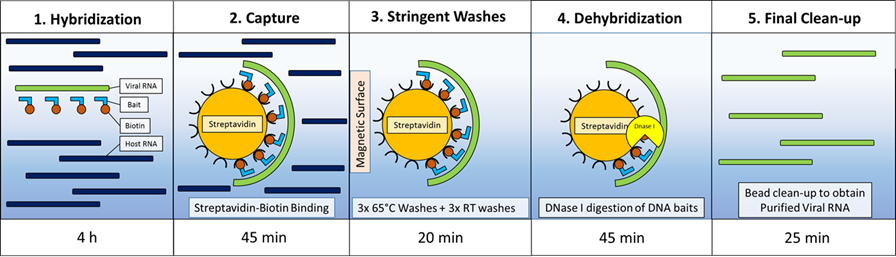

| Table S1: Reagents and volumes for hybridization step. | |
| --- | --- |
| **Components** | **Volume (uL)** |
| 2x Hybridization Buffer | 30 |
| Hybridization Component A | 12 |
| Diluted Baits | 4 |
| Nuclease-free water | 10 |

Table S2: Reagents and volumes for RT-qPCR.

| **Components** | **Volume (uL)** |
| --- | --- |
| RNA in Nuclease-free Water | 1.0 |
| 2x QuantiTect Probe RT-QPCR Master Mix Buffer | 5.0 |
| QuantiTect RT Mix | 0.1 |
| Nuclease-free Water | 1.65 |
| 10 µM Forward Primer | 0.9 |
| 10 µM Reverse Primer | 0.9 |
| 10 µM Probe | 0.45 |
| Total Volume | 10 |

Table S3: RT-qPCR program set on Applied Biosystems ViiA 7 Real Time-PCR System.

| **Sequence Name** | **Sequence (5’-3’)** |
| --- | --- |
| DENV1 F | ACTAGYGGTTAGAGGAGACC |
| DENV1 R | GGTCTCCWCTAACCTCTAGT |
| DENV1 Probe | FAM-ACCAGGGRAAGCTGTAYCYT-BHQ |
| DENV1 Standard | ACTAGTGGTTAGAGGAGACCCCTCCCGAAACACAACGCAGCAGCGGGGCCCAACACCAGGGGAAGCTGTACCCTGGTGGTAAGGACTAGAGGTTAGAGGAGACC |
| GAPDH F | GATTCCACCCATGGCAAATTC |
| GAPDH R | ATTTCCATTGATGACAAGC |
| GAPDH Probe | FAM-CGTTCTCAGCCTTGACGGTGCCA-BHQ |
| GAPDH Standard | TATGATTCCACCCATGGCAAATTCCATGGCACCGTCAAGGCTGAGAACGGGAAGCTTGTCATCAATGGAAATCCCATCA |

Table S4: Primers, probes and standards used with the respective modifications for RT-qPCR.

| **Program name** | **Temperature/°C** | **Time/min** | **Number of Cycles** |
| --- | --- | --- | --- |
| Reverse Transcription | 48 | 30 | 1 |
| PCR Initial Activation Step | 95 | 15 |  |
| Denaturation | 95 | 0.25 | 40 |
| Combined Annealing/Extension | 55 | 1 |  |

Table S5: Reagents and volumes used for polyadenylation of RNA samples.

| **Components** | **Volume (uL)** |
| --- | --- |
| **RNA** | 24 |
| **10x E. Coli Poly(A) Polymerase Reaction Buffer** | 3.2 |
| **ATP (10mM)** | 3.2 |
| **E. Coli Poly(A) Polymerase** | 1.6 |

Figure S1: DENV1 and GAPDH standard curves used for calculation of primer efficiency.

Standard Curve for DENV1

| <CT> | Log2(amount) |
| --- | --- |
| 31.1321 | 14.5567 |
| 25.4492 | 17.8787 |
| 20.7854 | 21.2006 |
| 15.8644 | 24.5225 |
| 11.7963 | 27.8445 |
| 8.5882 | 31.1664 |

Standard Curve for GAPDH

| <CT> | LOG2(copies/UL) |
| --- | --- |
| 32.5527 | 7.91289 |
| 29.2241 | 11.2348 |
| 25.4414 | 14.5567 |
| 21.7386 | 17.8787 |
| 18.1378 | 21.2006 |
| 14.5893 | 24.5225 |
| 11.1259 | 27.8445 |
| 8.23105 | 31.1664 |

Figure S2: Coverage plot against nucleotide position on DENV1 reference for pre-capture MinION sequencing run.

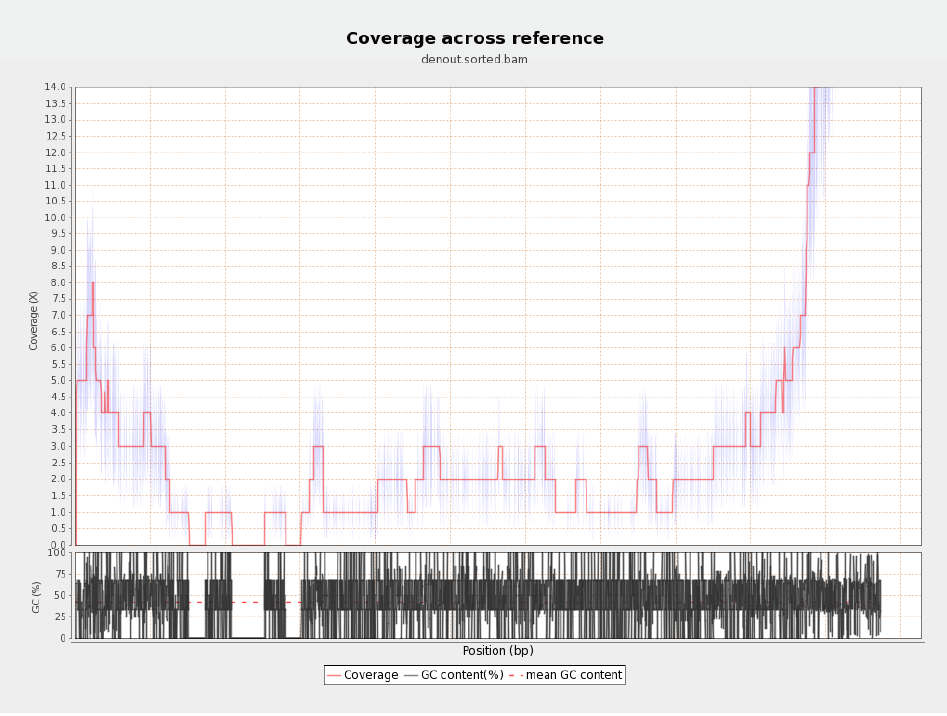

Figure S3: Coverage plot against nucleotide position on DENV1 reference for post-capture MinION sequencing run.

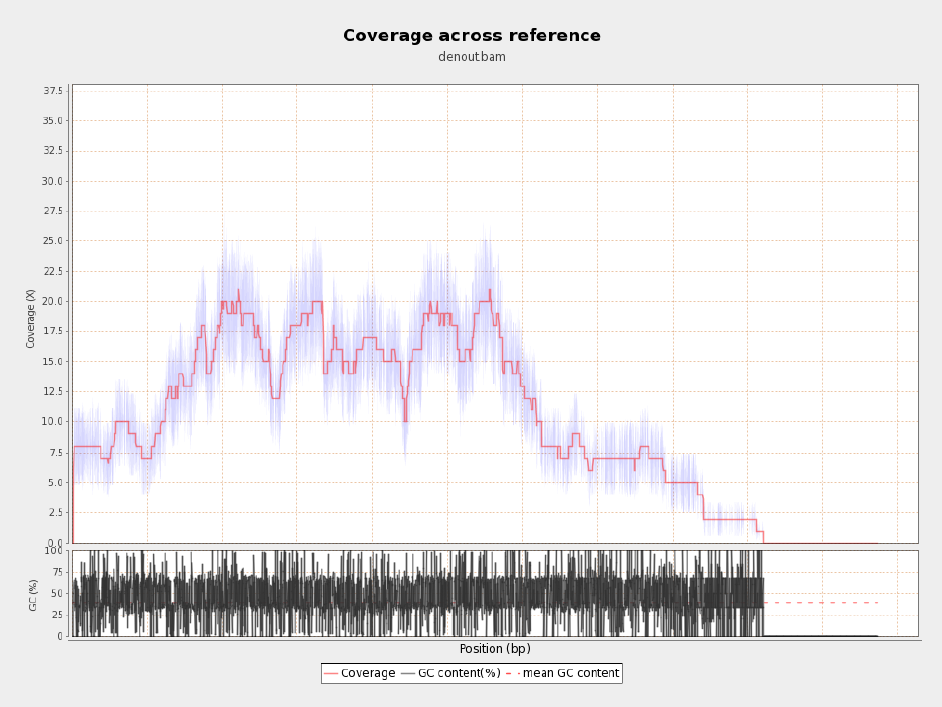

Figure S4: Coverage plot against nucleotide position on DENV1 reference for concentrated post-capture group MinION sequencing run.

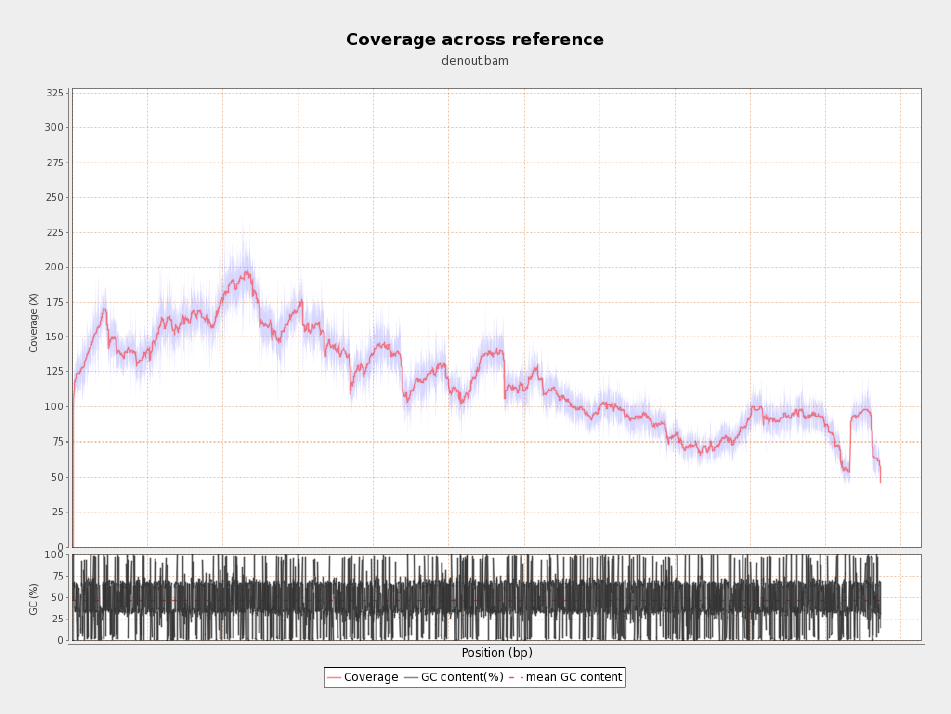
