## Supplementary material for "A Novel Method for the Capture-based Purification of Whole Viral Native RNA Genomes": All tables

Table 1: Comparison of absolute quantities of DENV1 and GAPDH used for calculation of purification factor.

| **Treatment Group** | **Primer** | **<C_q_>** | **Copies/µL** | **Total No. of Copies** | **Purification Factor** |
| --- | --- | --- | --- | --- | --- |
| Pre-capture | DENV1 | 10.47*±0.08* | 5.50x10^8^ | 8.25x10^9^ | 5.61x10^2^ |
|  | GAPDH | 14.75*±0.1* | 2.52x10^7^ | 3.78x10^8^ |  |
| Post-capture | DENV1 | 17.14*±0.09* | 1.92x10^7^ | 5.77x10^8^ |  |
|  | GAPDH | 29.63*±0.1* | 1.57x10^3^ | 4.72x10^4^ |  |
| Pre-capture | DENV1 | 12.13*±0.1* | 2.39x10^8^ | 3.58x10^9^ | 7.91x10^2^ |
|  | GAPDH | 17.04*±0.4* | 5.70x10^6^ | 8.55x10^7^ |  |
| Post-capture | DENV1 | 15.31*±0.1* | 4.83x10^7^ | 1.45x10^9^ |  |
|  | GAPDH | 29.75*±0.07* | 1.46x10^3^ | 4.37x10^4^ |  |

Table 2: Summary of bioinformatic analysis for pre and post-capture MinION sequencing runs.

|  | **Pre-capture** | **Post-capture** | **Concentrated post-capture** |
| --- | --- | --- | --- |
| **<DENV1 Coverage>** | *12.11* | *10.02* | *120.23* |
| **<DENV1 Mapping Quality>** | *34.31* | *31.60* | *36.99* |
| **DENV1 Error Read Rate (%)** | *16.49* | *17.61* | *15.27* |
| **Mapped to DENV1 (%)** | *0.63* | *31.35* | *77.47* |
| **Mapped to Human (%)** | *77.66* | *14.19* | *6.05* |
| **Percentage of DENV1 Genome Recovered ≥15X coverage (%)** | *7.05* | *27.45* | *99.91* |
| **Purification factor** | *272x* | | *1580x* |

Table 3: Summary of advantages and disadvantages of strategies for direct RNA sequencing of RNA viruses

| **Strategy Type** | **Employed in** | **Advantages** | **Disadvantages** |
| --- | --- | --- | --- |
| 1. Custom Adapter | Keller *et al.* 2018 | - Highest sequencing specificity - Rapid turnaround time | - Cannot sequence fragmented RNA - Foreknowledge of target sequence required |
| 2. Conventional Adapter | Wongsurawat *et al.* 2019  Pre-capture group | - Sequences fragmented RNA - No foreknowledge of target sequence - Rapid turnaround time | - Low sequencing specificity |
| 3. Conventional Adapter + Capture | post-capture group  concentrated post capture group | - Sequences fragmented RNA - Less foreknowledge of target sequence required - Higher sequencing specificity | - Increased turnaround time |
| 4. Conventional Adapter + host rRNA depletion | Wongsurawat *et al.* 2019 | - Sequences fragmented RNA - No foreknowledge of target sequence - Fast turnaround time | - Low sequencing specificity |
